# Deficiency of the Endocytic Protein *Hip1* Leads to Decreased *Gdpd3* expression, Low Phosphocholine, and Kypholordosis

**DOI:** 10.1101/383083

**Authors:** Ranjula Wijayatunge, Sam R. Holmstrom, Samantha B. Foley, Victoria E. Mgbemena, Varsha Bhargava, Gerardo Lopez Perez, Kelly McCrum, Theodora S. Ross

## Abstract

Deficiency of huntingtin interacting protein 1 (Hip1) results in degenerative phenotypes. Here we generated a *Hip1* deficiency allele where a floxed transcriptional stop-cassette and a human *HIP1* cDNA were knocked-in to intron 1 of mouse *Hip1* locus. *CMV-Cre-* mediated germline excision of the stop-cassette resulted in expression of HIP1 and rescue of the *Hip1* knockout phenotype. *Mxl-Cre-*-mediated excision led to HIP1 expression in spleen, kidney and liver, and also rescued the phenotype. In contrast, *GFAP-Cre-* mediated *HIP1* expression in brain did not rescue the phenotype. Metabolomics and microarrays of several *Hip1* knockout tissues identified low phosphocholine (PC) levels and low *Glycerophosphodiester Phosphodiesterase Domain Containing 3 (Gdpd3*) expression. Since Gdpd3 has lysophospholipase D activity that results in the formation of choline, a precursor of PC, *Gdpd3* downregulation could lead to the low PC levels. To test if *Gdpd3* contributes to the Hip1 deficiency phenotype, we generated *Gdpd3* knockout mice. Double knockout of *Gdpd3* and *Hip1* worsened the Hip1 phenotype. This suggests that Gdpd3 compensates for Hip1 loss. More detailed knowledge of how Hip1 deficiency leads to low PC will improve our understanding of HIP1 in choline metabolism in normal and disease states.

## INTRODUCTION

HIP1 was originally discovered as a protein that interacts with Huntingtin, the protein product of the gene mutated in Huntington’s disease (*1, 2*). HIP1 was later found to be an endocytic protein that binds clathrin, AP2 (*3-6*), inositol lipids (*7-9*) and actin (*10).* Knockout of *Hip1* alone (*11, 12*) or with its only known mammalian relative, Hip1-related (*Hip1r) (13, 14*), results in a degenerative phenotype. This phenotype, which is more severe in double-knockout mice, includes testicular degeneration, spinal defects, and lowered body weight. Although it has been shown that endocytosis of AMPA receptors in neurons is disrupted in knockout mice *(11)*, mechanisms for most of the knockout phenotypes are unknown as are the roles of HIP1 in normal and neoplastic tissues.

HIP1 was first linked to receptor tyrosine kinase (RTK) signaling when it was identified as a chromosomal translocation partner with the platelet-derived growth factor β receptor (*PDGFβR*) gene in leukemia (15). This *HIP1/PDGFβR* translocation is a member of a large family of chromosomal translocations involving the *PDGFβR* gene (*16-19).* Recently, others have identified *HIP1* as a partner in chromosomal translocations involving the anaplastic lymphoma kinase (*ALK*) gene in non-small cell lung cancer (20-22). Like *HIP1/PDGFβR*, the *HIP1/ALK* translocation is a member of a large family of chromosomal translocations that involve the *ALK* gene.

In addition to the *HIP1/PDGFβR* and *HIP1/ALK* mutations, the HIP1 protein itself is expressed at high levels in several other cancers (*23-25*) and overexpression of HIP1 transforms fibroblasts (24) and prostate epithelial cells (26). This transforming activity is also linked to RTKs, as HIP1-transformed cells display increased levels of EGF receptor (EGFR) (*24).* Indeed, HIP1 prolongs the half-lives of RTKs, such as EGFR and PDGFβR (*8).* Additionally, treatment of the HIP1-transformed cells with an EGFR inhibitor reverses the transformed phenotype (24). One possible explanation for EGFR overexpression in these cancers is that HIP1-dependent stabilization of RTKs occurs via clathrin or membrane vesicle sequestration. Low levels of clathrin or altered membrane dynamics could decrease endocytosis-mediated RTK degradation. This would then prolong RTK signaling and promote transformation.

Here, we generated *Hip1* deficient mice with the option to conditionally express a single copy of human *HIP1* in specific tissues to investigate how *Hip1* deficiency leads to degeneration. With this new model, we discovered that the brain does not require Hip1 for normal physiology, but that Hip1 expression in spleen, liver, and kidney may be important for normal physiology. We also assayed a variety of *Hip1* deficient tissues for alterations in metabolism and gene expression and discovered that low phosphocholine (PC) levels and *Gdpd3* expression were associated with *Hip1* deficiency. We also observed *Gdpd3/Hip1* double deficient mice to have spinal degeneration at younger ages compared to single *Hip1* deficient mice suggesting Gdpd3 partially compensates for the Hip1 deficiency. Low PC levels in Hip1 deficient tissue is opposite to elevated levels of HIP1 and PC found in cancers (27).Thus, HIP1 may, via endocytosis of lipids or via effects on gene expression, lead to normal or abnormal levels of choline-related metabolites in both healthy and diseased tissues.

## MATERIALS AND METHODS

### Generation of *Hip1* deficient mice and conditional *HIP1* knockin mice

The targeting vector was constructed to generate a knock-in mouse allele that conditionally expresses human HIP1 and enhanced GFP (EGFP; **Figure 1A**). The objective of this project has been to create a knock-in mouse model conditionally expressing human Hip1 and eGFP using homologous recombination in mouse C57BL/6 embryonic stem cells and subsequent blastocyst injection of the appropriate targeted ES cells to create the gene targeted mice.

The mouse *Hip1* genomic DNA sequence was retrieved from mouse chromosome 5 from the Ensembl database and used as reference in this project. The mouse BAC DNA containing *Hip1* gene was used as templates for generating the homology arms and the southern probes for screening targeted events. The 5’ homology arm (~5.4 kb, LHA) and the 3’ homology arm (~3.6 kb, SHA) were generated by PCR using high fidelity Taq DNA polymerase. The Lox-Stop-Lox cassette (~1.5 kb, LSL) was amplified from LSL-pZero-2 vector. These fragments were cloned in the FtNwCD or pCR4.0 vector, and were confirmed by restriction digestion and sequencing. The hHip1 cDNA (~3 kb) was amplified from pcDNA3-hHip1 plasmid. The IRES (~0.6 kb) and eGFP (~0.9 kb) sequences were amplified from pIRES2-AcGFP1 and pEGFP-N1 plasmid, respectively.

The final vector was obtained by standard molecular cloning. Aside from the homology arms, the final vector also contains a Lox-Stop-Lox cassette (~1.5 kb), mHip1 partial intron1+ partial exon 2, hHip1 cDNA, IRES, EGFP + polyA, Frt sequences flanking the Neo expression cassette (the neo cassette was used for positive selection of the electroporated ES cells), and a DTA expression cassette (for negative selection of the ES cells). Unique restriction sites SacII and PacI were added to flank the hHip1 cDNA for easy swapping in mutant cDNA in the future. The final vector was confirmed by both restriction digestion and sequencing analysis. NotI was used for linearizing the final vector prior to electroporation.

### ES cell gene targeting

Thirty μg of NotI-linearized final gene targeting vector DNA was electroporated into ~10^7^ C57BL/6 ES cells and selected with 200 μg /ml G418. Two plates of G418 resistant ES clones (~192) were selected for screening. The primary ES screening was performed with 3’ PCR. Approx. 40 potential targeted clones were identified from one plate. Six clones (A8, B4, C7, C11, E6 and F4) were expanded for further analysis.

Upon completion of the ES clone expansion, additional Southern confirmation analysis was performed. Based on this analysis, five out of the six expanded clones (B4, C7, C11, E6 and F4) were confirmed for homologous recombination with single neo integration. Clones C7 and C11 were then transfected with a Flp expressing plasmid. Eight G418 sensitive clones (C7/A1, C7/A10, C7/B7, C11/B5, C11/B9, C11/C8, C11/D7 and C11/E8) were identified and confirmed as Neo deleted clones by PCR.

### Southern blot

The 5’ and 3’ external probes were generated by PCR and were tested by genomic Southern blot analysis for screening of the ES cells. The probes were cloned in the pCR4.0 backbone and confirmed by sequencing. Southern blot to distinguish the targeted *Hip1^LSL^* allele and the wild-type allele was performed with both probes as described (*12*).

### Generation of *Gdpd3* deficient mice

The targeting strategy for the generation of the *Gdpd3* knockout allele was based on NCBI transcript NC_000073_NM_024228_2(Figure 5A). The constitutive *Gdpd3* knock out mouse was generated with the assistance of Taconic biosciences and CRISPR/Cas9-mediated genome editing. Of all potential sgRNAs, two were selected for their position and minimal potential off-target effects. Guide RNAs target sequences before exon 2 (proximal sgRNA, 5’-GAG AGG CCA GTT CAA CTG AT-3’) and after exon 6 (distal sgRNA, 5’GAC GGG GAT TCG ACA ATT GC-3’). The resulting alteration via Non-Homologous End Joining (NHEJ) deleted the functionally critical exons (2-7), generated a frame-shift mutation from exon1 to all downstream exons and introduced a premature Stop codon in exon 7. In addition, the resulting transcript was predicted to be a target for non-sense mediated RNA decay and therefore not expressed at a significant level. The guide RNAs and Cas9 mRNA/protein were injected into single celled zygotes. C57BL/6J mice were used to generate these mice. Mice were tail genotyped using the following primers:

Oligo-1 5’-TCC ATG TAG GGT GGA GTG AGC-3’

Oligo-2 5’-AAG GGG CTC GTA GGG GAA G-3’

Oligo-3 5’-ACA CAG GAA GGA CCA GGG C-3’

Amplicon sizes for the wild type allele was 501 bp (with oligos 1 and 3) and for the deletion allele was 295 bp (with oligos 1 and 2). These PCR products were cloned and sequenced. Indel modifications were distinguished from unmodified wildtype sequences by heteroduplex analysis via capillary electrophoresis.

### Mice

CMV-Cre;Hip1^LSL/LSL^, *Mx1-Cre;Hip1^LSLlLSL^* or *GFAP-Cre; Hip1^LSL/LSL^* mice were generated by crossing the *Hip1^LSL/LSL^* mice with mice transgenic for the *CMV-Cre* (28), *Mx1-Cre (29*) or the *GFAP-Cre* genes (30) respectively. Expression of Cre recombinase from the *Mx1-Cre* transgene was induced by intraperitoneal injection of mice with 250μg pIpC (Sigma, St. Louis, MO) in 100μl PBS at 2-day intervals for 2 weeks. Mice were housed in the Unit for Laboratory Animal Medicine at the University of Texas Southwestern Medical School under specific pathogen-free conditions. All mouse experiments were conducted after approval of the UT Southwestern Medical Center Committee on the Use and Care of Animals.

### Genotyping

The *Hip1^LSL^* conditional allele and the *Gdpd3* knockout allele were genotyped from tail snips using real-time PCR assays designed by Transnetyx. For Hip1, assays were designed to detect wild-type *Hip1* and the Cre-recombined (excised LSL cassette) and un-recombined mutant alleles. For *Gdpd3*, an assay was designed to detect wild-type *Gdpd3* allele.

### Western blot

Tissues and cells were extracted in lysis buffer containing protease and phosphatase inhibitors (50 mM Tris pH 7.4, 150 mM NaCl, 1% Triton X-100, protease inhibitors (Roche), 30 mM sodium pyrophosphate, 50 mM NaF, 100 μM sodium orthovanadate), cleared of unbroken cells by centrifugation, and diluted to a protein concentration of 0.5 to 2.0 mg/mL. Whole cell lysates in Laemmli buffer were separated on 6% or 10% SDS-PAGE gels and transferred to nitrocellulose membranes. Membranes were probed with the following antibodies: Mouse monoclonal anti-human HIP1 4B10 (1:1,000) or anti-mouse Hip1 1B11 (1:1,000) and rabbit polyclonal anti-human HIP1 UM410 and UM323 (1:5,000). Blots were then incubated with horseradish peroxidase (HRP)-conjugated mouse or rabbit secondary antibodies (1:10,000; GE healthcare) and developed using chemiluminescence (Pierce).

### Metabolomics

Mouse tissues from age matched *Hip1* deficient (n=7) and *HIP1* rescued(n=7) or *Gdpd3^+I+^* (n=5) and *Gdpd3^-/-^* (n=5) mice were rapidly removed (within 2 minutes of sacrifice) and snap frozen. *Hip1* deficient mice were <3 months of age. *Gdpd3* knockout mice were 2–6 months of age. Frozen tissues (~100mg each) were homogenized in ice-cold 80% methanol. Following a vigorous vortex, cell debris was removed by centrifugation. The supernatant was evaporated with a SpeedVac to generate a metabolite pellet which was reconstituted in 0.03% formic acid in analytical-grade water before liquid chromatography-tandem mass spectrometry (LC-MS/MS) analysis. LC-MS/MS was performed as described previously (*31*).

To gain an initial overview of the results, a multivariate principle component analysis of all 120 metabolites that were measured was performed. All analyses were carried out using SIMCA-P (version 13.0.1; Umetrics). Although there were some differences between the two genotypes, the main changes were in choline metabolites as described in the results. The Variable Importance in Projection (VIP) values were calculated with the R statistical package. All other statistical analyses of the mass spectrometry data were performed with Prism 5.0c GraphPad Software. P values were calculated with unpaired Student’s t-test.

### Microarray

Total RNA was prepared from various tissues using TRIzol (Invitrogen) extraction and treated with DNAse (QIAGEN) to remove genomic DNA contamination. RNA from four (two 1-3 and two 5-6 month old) mice from each genotype (*Hip1^LSL/LSL^, Hip1^HIP1/HIP1^* or *Hip1* wild type) was used. Microarray was carried out using Illumina MouseWG-6 V2 Beadchip whole-genome expression array (Illumina, Inc.). Each RNA sample was amplified using the Ambion TotalPrep RNA amplification kit with biotin UTP (Enzo) labeling. The Ambion Illumina RNA amplification kit used T7 oligo(dT) primer to generate single stranded cDNA followed by a second strand synthesis to generate double stranded cDNA which was then column purified. In vitro transcription with T7 RNA polymerase generated biotin-labeled cRNA. The cRNA was then column purified, checked for size and yield using the Bio-Rad Experion system, and 1.5ug of cRNA was hybridized to each array using standard Illumina protocols.

Streptavidin-Cy3 (Amersham) was used for detection. Slides were scanned and fluorescence intensity captured with Illumina HiScan. Expression values were extracted using GenomeStudio2010.2. The data was background subtracted and quantil-normalized using the MBCB algorithm (*32-34).*

### Real Time PCR for *Gdpd3* Expression

Total RNA (2 μg) was used for reverse transcription using iScript Advanced cDNA Synthesis Kit (BIORAD) according to manufacturer’s instructions. Real Time PCR was then performed using StepOnePlus Real-Time PCR system (Applied Biosystems) with Taqman gene expression master mix and specific Taqman gene expression assays [mGdpd3 (Ms00470321_m1), mGapdh (Ms99999915_g1); Life Technologies]. Each sample was measured in triplicate, and all data were normalized to the housekeeping gene *Gapdh* as a control for RNA quantity and sample processing.

## RESULTS

### Generation of a knock-in allele of human HIP1

Here, we report the generation of a novel *Hip1* knockout allele (*Hip1^LSL^;* Figure 1A). This allele also has the human *HIP1* cDNA knocked into the mouse *Hip1* locus downstream of a *loxP*-flanked transcriptional stop cassette (LSL). Having previously found that a multi-copy-transgenic *HIP1* cDNA allele was able to rescue the *Hip1* and *Hip1r* double knockout degenerative phenotype (13), we hypothesized that *HIP1* as a single copy allele may also substitute for *Hip1* (or *Hip1r*) deficiency phenotypes. This conditional knock-in allele allows for use of specific *Cre* alleles to guide tissue-specific HIP1 expression together with the endogenous mouse *Hip1* promoter to further regulate expression. In addition to the conditional nature of this allele that allows for expression of HIP1 in specific tissues at specific times, there were other potential advantages of this targeting strategy. First, since HIP1 is expressed at high levels in many cancers (*24, 25, 35, 36*), humanization of a mouse allele could be of future use for preclinical *in vivo* HIP1 targeted drug testing. Second, this type of humanization addresses the question of whether the many introns of the large 250 Kb *Hip1* locus are required for regulation of the *Hip1* gene. We reasoned that if the expression of *HIP1* from a single knocked-in copy does not restore normalcy to the *Hip1* deficient mice, the complex genomic structure of the allele is necessary for HIP1 expression.

We successfully targeted ES cells (**Figure 1B**) and generated three independent mouse lines carrying the *Hip1^LSL^* allele (the “knockout” allele). Homozygous and heterozygous knockout mice were born at predicted Mendelian frequencies. To generate the germline “humanized” allele, *Hip1^LSL/+^* mice were crossed with transgenic *CMV-Cre* mice (28). The CMV promoter is activated in all tissues including germ cells. The resulting *Hip1^Hip1^* humanized allele was used to generate fully humanized, *Hip1^Hip1/Hip1^* mice (CMV-Cre allele was crossed out). *Hip1^Hip1/Hip1^* mice could be distinguished from *Hip1* knockout (*Hip1^LSL/LSL^*) mice by GFP expression in peripheral blood (**Figure 1C**) or by the presence of human HIP1, rather than mouse Hip1 protein, in mouse tissues (**Figure 1D, bottom**).

**Figure 1.**
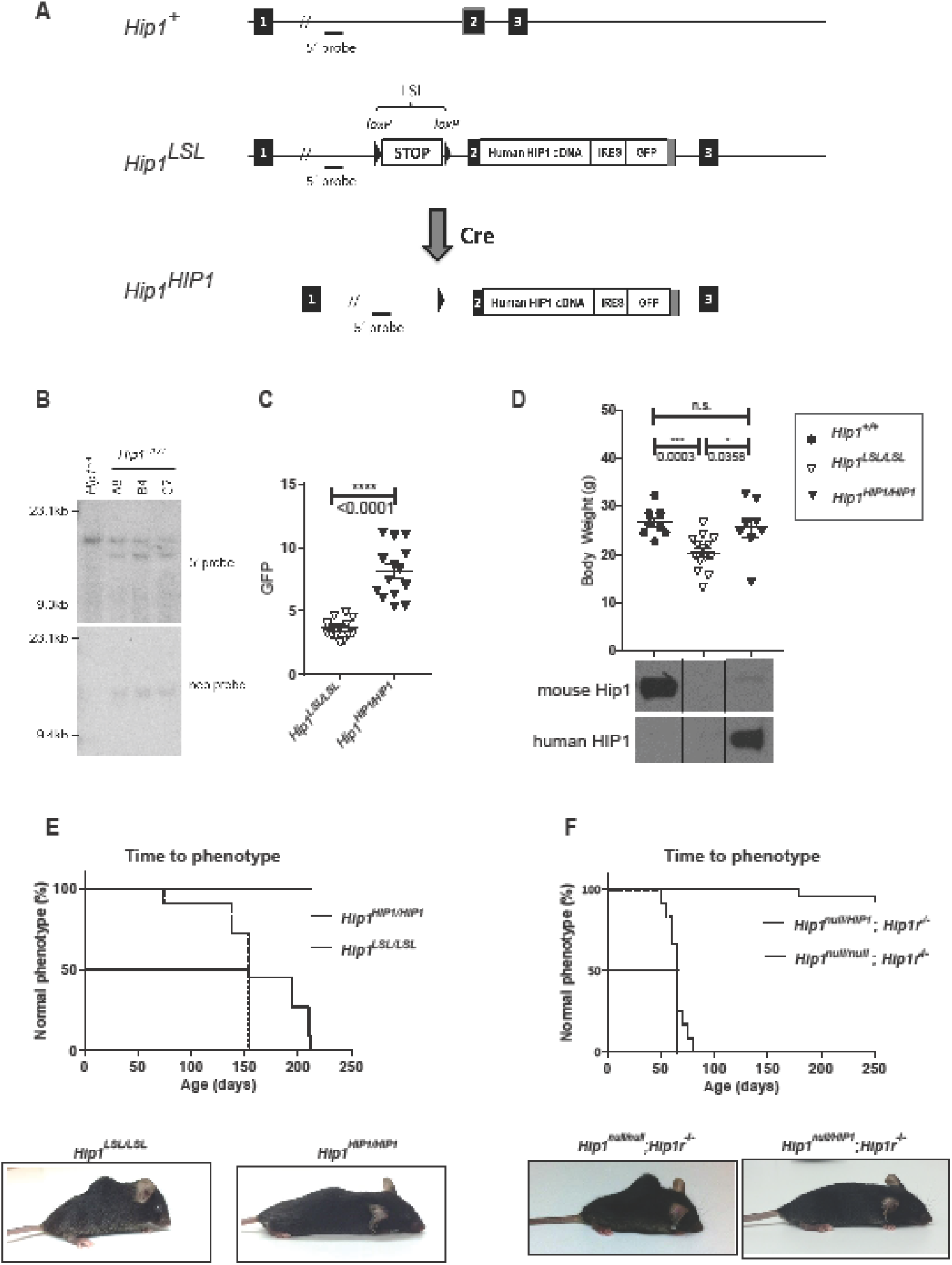
Generation of *Hip1* Deficient and Conditionally *HIP1* Humanized Mice. **(A)** Schematic of the first 3 of the 32 exon murine *Hip1* locus (*Hip1^+^*), the targeted knock-in allele with a floxed stop cassette (LSL) and a human *HIP1* cDNA inserted into the murine *Hip1* locus by homologous recombination (*Hip1^LSL^*), and the Cre-mediated recombined allele with the excised stop cassette (*Hip1^HIP1^*). The following features are indicated: stop cassette bracketed by *loxP* recombination sequences (LSL), partial mouse exon 2 fused to human *HIP1* cDNA-IRES-EGFP poly A tail (light grey box) and genomic hybridization probes. **(B)** Southern blot confirmation of successfully targeted ES cell lines. Shown are wild-type (*Hip1^+/+^*) and ES cell lines carrying *Hip1^LSL^* targeted allele (A8, B4, and C7). Genomic DNA was digested with EcoRl or HindlII, and southern blotted with either the 5’ probe to yield a 15.7 kb band corresponding to the wild-type allele and 13.4 kb band corresponding to the recombined allele or the 3’ neomycin probe to yield a 15.5 kb band corresponding to the recombined allele, respectively. The neomycin expression cassette was then excised by transfection of the cells with a Flp expressing plasmid. **(C)** A *CMV-Cre* transgene was used to generate a germline Cre-mediated recombined allele, *Hip1^HIP1^*, as delineated in **part A**. Here GFP expression in peripheral blood WBCs was quantitated to confirm that excision of stop cassette leads to GFP expression in *Hip1^HIP1/HIP1^* mouse tissue. Data represent mean +/- S.E.M., n=14 per group. **(D) (Top)** Expression of human HIP1 from the *Hip1* locus prevents the weight loss of *Hip1* deficient mice. Data represent mean +/- S.E.M. n=8-16. ***p < 0.005, *p < 0.05, n.s., not significant. Ages ranged between 5 and 6.5 months. (Bottom) Western blot analysis confirmed that expression of human HIP1 protein replaces mouse Hip1 protein in lung tissue from *Hip1^HIPI/HIPI^* mice. The top panel represents western blot with a human specific polyclonal antibody (UM323) and the bottom panel a mouse specific monoclonal antibody (UM.1B11). (E) Expression of human HTP1 from the *Hip1* locus prevents kypholordotic and weight loss phenotypes in all mice. **(Top)** The Kaplan-Meier curves depict the percentage of mice with a normal phenotype as a function of time in days since birth. *Hip1* deficient mice (*Hip1^LSL/LSL^*) are represented by the dotted line and the fully humanized HTP1 mice (*Hip1^HIPI/HIPI^*) are represented by the solid line. n=11 per genotype. **(Bottom)** Representative mice at 6 months of age with and without the *Hip1* deficiency associated kypholordosis and diminished weight. Note that the HTP1 humanized mouse (*Hip1^HIP1/HIP1^*) displays a slight kyphosis that is also observed in wild type mice. Without the lordosis displayed by the *HipI^LSL/LSL^* mouse, this slight kyphosis is normal. **(F)** Kypholordic phenotype of Hip1 and Hip1r double knockout mice is rescued by a single copy of human HTP1. **(Top)** The Kaplan-Meier curves depict the percentage of mice with a normal phenotype as a function of time in days since birth. Rescued *Hip1^null/HIP1^; Hip1r^-/-^* mice are represented by the solid line and double knockout *HipI^null/null^*; *Hip1r^-/-^* mice by the dotted line. (**Bottom**) Representative photographs of 2 month old *Hip1^null/null^; Hip1r^-/-^* mouse with kypholordosis and an age-and gender-matched *HipI^null/HIP1^; Hip1r^-/-^* rescue mouse with no phenotype.

### Rescue of the *Hip1* deficiency phenotype by a knocked-in human *HIP1* cDNA

Although indistinguishable at birth from their wild type or heterozygous littermates, by six months of age, all *Hip1* knockout mice (*Hip1^LSL/LSL^*) were afflicted with lowered body weight and severe spinal defects. In contrast, the *HIP1* humanized mice (*Hip1^Hip1/Hip1^*), remained normal in weight (**Figure 1D, top**) and were without a kypholordotic spine (**Figures 1E**). These data indicate that a single copy of human HIP1 cDNA is able to fully substitute for the mouse *Hip1* gene. This was also supported by the ability of the *Hip1^Hip1^* allele to fully rescue the *Hip1/Hip1r* double knockout phenotype, which was more severe than in the single knockout mice (**Figures 1F**).

### HIP1 expression in the brain does not prevent the *Hip1* knockout phenotype

Because *Hip1* is expressed at high levels in the brain and the Hip1 protein interacts with Huntingtin, the product of the gene mutated in Huntington’s syndrome (*1*, 2), we next asked if restricted expression of HIP1 in neural tissues would rescue the *Hip1* deficiency phenotypes. Because Hip1 has also been shown to be required for AMPA receptor trafficking in neurons (*11*), we predicted HIP1 expression in the brain would rescue the *Hip1* deficiency phenotypes. To restrict HIP1 expression to neural tissue, we crossed the *Hip1^LSL/LSL^* mice with transgenic *GFAP-Cre* mice (*30).* The *GFAP* promoter in these mice is active in most glial and neuronal cells of the brain and spinal cord. As expected, western blot analysis of human HIP1 expression in *GFAP-Cre*; *Hip1^LSL^* ^*/LSL*^ brains demonstrated levels similar to the germline humanized *Hip1^HIP1/HIP1^* mouse brains and HIP1 protein was not detected in spleen, lung, kidney or liver (**Figure 2A**). To our surprise, the degenerative phenotypes associated with *Hip1* deficiency were *not* rescued by *GFAP-Cre* mediated expression of HIP1 (**Figure 2B**). These *GFAP-Cre* data suggest *Hip1* deficiency in the brain does not contribute to the *Hip1* deficiency phenotype.

**Figure 2.**
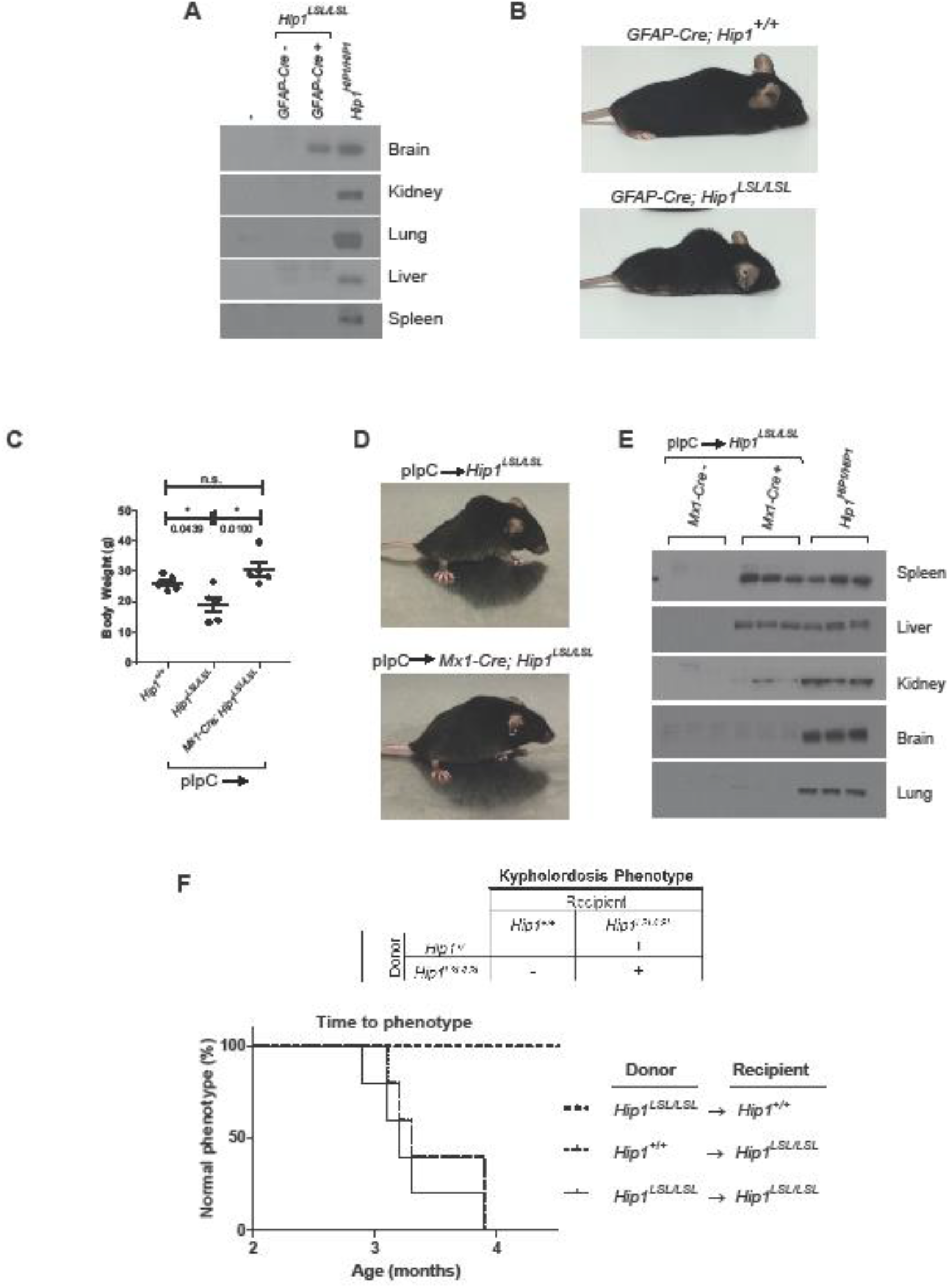
Tissue-specific Rescue of *Hip1* Deficiency with Human HIP1. **(A)** Western blot for tissue-specific expression analysis of human HTP1 in brain from *GFAP-Cre; HipI^LSLlLSL^* mice. Tissue extracts from representatives of each genotype were analyzed. The human specific HTP1 antibody UM323 was used to detect HTP1 in the brain and showed no detectable HIP1 in spleen, liver, kidney, and lung tissues. **(B)** The *Hip1* deficiency phenotype was not rescued by expression of human HIP1 in the nervous system with the *GFAP-Cre* transgene. Representative photographs of 6 month old *GFAP-Cre;Hip1^LSL/LSL^* and *GFAP-Cre;Hip1^+/+^* mice. **(C)** At 6 weeks of age, *Hip1^LSL/LSL^* and *Mxl-cre*; *Hip1^LSL/LSL^* mice were treated with pIpC to induce *Mxl-Cre* mediated expression of human HIP1 in the hematopoietic system, kidney and liver. Expression of *Mx1-Cre* prevented the weight loss observed in *Hip1* deficient mice (*Hip1^LSL/LSL^*). All mice were between 5.5 and 6 months of age as this is when the weight loss was most apparent. Data represent mean +/- S.E.M. n=5. n.s., not significant, *p < 0.05 **(D)** The kypholordotic spinal curvature in *Hip1* deficient mice is rescued by *Mxl-Cre* mediated expression of human *HIP1.* Representative photographs of 4 month old pIpC treated *Hip1^LSL/LSL^* and *Mxl-cre*; *Hip1^LSL/LSL^* mice. **(E)** Western blot for tissue-specific expression analysis of human HIP1 in spleen, liver and kidney tissues. Tissue extracts from three mice of each genotype were analyzed. The human specific HIP1 antibody UM323 was used to detect HIP1 in the spleen, liver and kidney with no detectable HIP1 in brain and lung tissues. **(F)** Kaplan-Meier curve depicting the lack of or the progression of the development of the Kypholordotic phenotype in bone marrow transplanted mice. Transplantation of Hip1 deficient (*Hip1^LSL/LSL^*) bone marrow into irradiated wildtype (*Hip1^+/+^*) mice does not lead to the development of the Kypholordotic phenotype. Transplantation of wildtype (*Hip1^+/+^*) bone marrow into irradiated Hip1 deficient (*Hip1^LSL/LSL^*) mice does not prevent the development of the Kypholordotic phenotype. n=5 per group.

### *Mxl-Cre* mediated HIP1 expression rescues *Hip1* deficient mice

Next we crossed *Hip1^LSL/LSL^* mice to *Mx1-Cre* transgenic mice to express HIP1 in adult liver, kidney, and the hematopoietic system (29). We did this because these tissues express high Hip1 levels and the *Hip1* deficiency phenotype develops only in adulthood. The *Mx1* promoter is activated by interferon, which can also be induced by the synthetic double-stranded RNA, polyinosinic-polycytidylic acid (pIpC) (*29*). Mice were induced with pIpC at 6 weeks of age prior to the onset of the degenerative phenotype. The induced *Mx1-Cre;Hip1^LSLlLSL^* mice (n=5) were of normal weight (**Figure 2C**) and without spinal defects at 4 months of age (**Figure 2D**), whereas induced *Hip1^LSL /LSL^* littermates were severely affected with the degenerative phenotype.

The pIpC induced *Hip1^LSLlLSL^* mice and *Mx1-Cre;Hip1^LSLlLSL^* rescued mice were evaluated for human HIP1 expression to confirm specificity of the *Cre* allele. Western blot analyses for human HIP1 protein demonstrated that pIpC-induced recombination occurred most effectively in spleen and liver. Kidney did not show as robust HIP1 expression. As expected, HIP1 was not detected in brain or lung tissues (**Figure 2E**). These *Mx1-Cre* data suggest that *Hip1* deficiency in adult liver and/or hematopoietic tissues (and less likely kidney) contributes to development of *Hip1* deficiency phenotypes.

### Bone marrow transplant of wild type bone marrow does not prevent the *Hip1* knockout phenotype

We transplanted wildtype bone marrow into *Hip1* deficient mice to test if replacement of the *Hip1* deficient bone marrow with normal bone marrow could rescue the kypholordotic phenotype. Bone marrow cells from 6-week-old wildtype (CD45.1^+^) or *Hip1^LSLlLSL^* mice were transplanted into 6-week-old lethally irradiated *Hip1^LSLlLSL^* mice (CD45.2^+^) (n=5 per group). These mice all developed a kypholordotic phenotype indicating that irradiation, wild type bone marrow (confirmed by the replacement of the CD45.2^+^ cells by CD45.1^+^ cells), or the transplant process in general does not prevent the phenotype (Figure 2F).

We also transplanted *Hip1* deficient bone marrow into wild type mice to test if Hip1 deficiency in bone marrow can cause the development of the *Hip1* mutant kypholordotic phenotype. Whole bone marrow cells from 6-week-old *Hip1^LSL/LSL^* CD45.2^+^) mice were transplanted into 6-week-old lethally irradiated wildtype (CD45.1^+^) mice (n=18). These mice did not develop the phenotype despite living to one year of age. Replacement of wildtype bone marrow with *Hip1* deficient bone marrow was confirmed by replacement of CD45.1^+^ cells by CD45.2^+^ cells (**data not shown**). In sum, Hip1 deficiency in bone marrow is insufficient for the development of the phenotype and Hip1 expression in the bone marrow is insufficient for the rescue of the phenotype.

### *Hip1* deficient tissues have low phosphocholine (PC) levels

The adult onset weight loss, generalized weakness and spinal defects of *Hip1* deficient mice raise the possibility of a progressive metabolic defect. Prior investigations to identify abnormalities in specific *Hip1* deficient cell types, such as osteoblasts or osteoclasts, or to identify endocrine abnormalities (e.g. altered thyroid or other hormone levels) were tested and eliminated (*13*). We therefore used mass spectrometry to compare levels of >100 metabolites in wildtype versus *Hip1* deficient tissues. Deletion of *Hip1* resulted in altered concentrations of multiple choline related metabolites, although choline levels were not changed in any tissues tested (brain, kidney, lung, liver, spleen). The most striking change was in phosphocholine (PC) which is generated by phosphorylation of choline by choline kinase. PC was significantly decreased in all *Hip1*-deficient tissues tested, except brain where it still trended low (**Figure 3A, B**). The largest observed decrease in PC was in the liver (by 0.41-fold, p=0.0003, VIP=1.8343), a tissue where choline metabolism is particularly relevant (Corbin and Zeisal, 2012). Because PC is a sentinel choline metabolite that reflects overall phosphatidylcholine pathway homeostasis (37), these data suggest a general change in choline metabolism that results from loss of *Hip1* expression. That choline metabolism is perturbed in *Hip1* deficient tissues was further supported by additional alterations observed in acetylcholine, betaine aldehyde, trimethylglycine/betaine and dimethylglycine (**DATASET S1**). However, no changes were detected in lysophosphatidylcholine or glycerophosphocholine.

**Figure 3.**
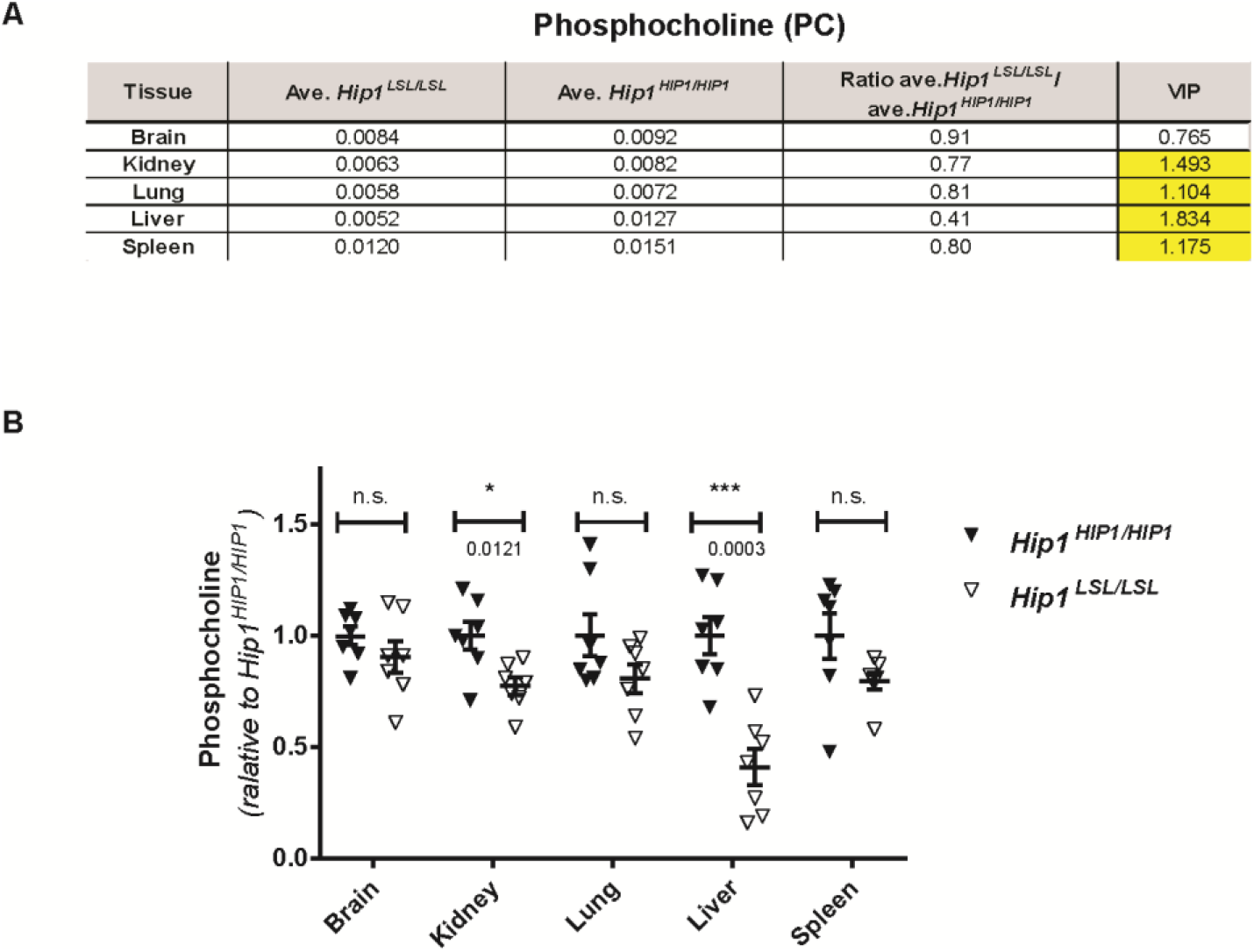
*Hip1* Deficient Mice have Low Phosphocholine Levels. **(A)** Of the 120 metabolites that were measured with LC-MSlMS, the most robust metabolic change was decreased phosphocholine (PC) levels. All tissues except brain from *HipI* deficient mice (*Hip1^LSL/LSL^*) showed significantly decreased PC levels compared to HTP1 rescued mice (*Hip1^HIP1/HIP1^*). The units are peak areas normalized to total ion current (a marker of total metabolite abundance).The VTP or Variable Tmportance in Projection number is a measure of statistical significance. When it is above 1.0, the metabolite change is considered statistically significant. **(B)** The same data in part A is shown as dot plots to visualize variability between samples. Data is expressed as relative to the average of levels in *HIP1* rescued tissues (*Hip1^HIP1/HIP1^*). Data represent mean +/- S.E.M., n=7 per group. n.s., not significant, *p < 0.05, ***p < 0.0001

There were other intriguing metabolite changes in *Hip1* deficient tissues (**DATASET S3**). One robust change was increased homocysteine in several *Hip1* deficient tissues. Of the tissues tested, homocysteine was increased the most in the liver (13.41-fold, p=0.002, VIP=1.8; **DATASET S2**). The concomitant decreases in pyridoxamine (a vitamer of vitamin B6) and cystathionine suggested that the increased homocysteine could be a result of a defective trans-sulfation pathway (pyridoxamine aids conversion of homocysteine to cysteine which is then converted to cystathionine). However, vitamin B6 levels were normal in the serum of *Hip1* deficient mice (**data not shown**). Increased homocysteine could also be due to an altered methionine cycle. In fact, S-adenosylhomocysteine (SAH) levels were increased and S-adenosyl methionine (SAM) levels were decreased in several tissues of *Hip1* deficient mice (**DATASET S2**), suggesting changes in steady state methylation. However, we did not detect changes in histone methylation by western blot analysis in tissues or by mass spectrometry (data not shown). Another possibility is that increased homocysteine was simply due to altered choline metabolism. Choline is metabolized to betaine, which acts as a methyl donor in the conversion of homocysteine to methionine. Since the liver showed the greatest decreases in PC and acetylcholine and the greatest increase in homocysteine of all tissues tested, this hypothesis is plausible. It is possible that high homocysteine redirects choline for betaine synthesis at the expense of acetylcholine and PC in order to maintain methionine homeostasis. In sum, these data suggest that Hip1 deficiency results in altered choline metabolism that includes elevated homocysteine levels.

### *Hip1* deficient mice lose *Gdpd3* expression

In order to identify potential mediators of the Hip1 deficiency phenotypes and the altered metabolome, we compared gene expression profiles from tissue of *Hip1^+/+^, Hip1^LSL/LSL^*, and *Hip1^HIP1/HIP1^* mice by microarray analysis. As expected, mouse *Hip1* was not expressed in either *Hip1^LSL/LSL^* knockout or *Hip1^HIP1/HIP1^* rescued mice but was expressed in *Hip1^+/+^* wildtype mice (data not shown). We identified *Gdpd3* (also known as *Gde7*) as a gene with a robust and reproducible decrease in expression in all tested tissues (**DATASET S4**). *Gdpd3 is* a gene whose protein product is a member of a glycerophosphodiester phosphodiesterase domain (Gdpd) family of enzymes, also known as the Gde family (*38, 39*). Expression patterns of the six other known members of the Gdpd family were not changed in *Hip1* deficient mice (**Figure 4A**). These decreases of *Gdpd3* expression were confirmed with qPCR (**Figure 4B**). Decreased expression of *Gdpd3* is not a consequence of the kypholordic phenotype since young, phenotypically “normal” Hip1 deficient mice show reduced *Gdpd3* expression levels similarly to old, kypholordotic mice (**data not shown**).

**Figure 4.**
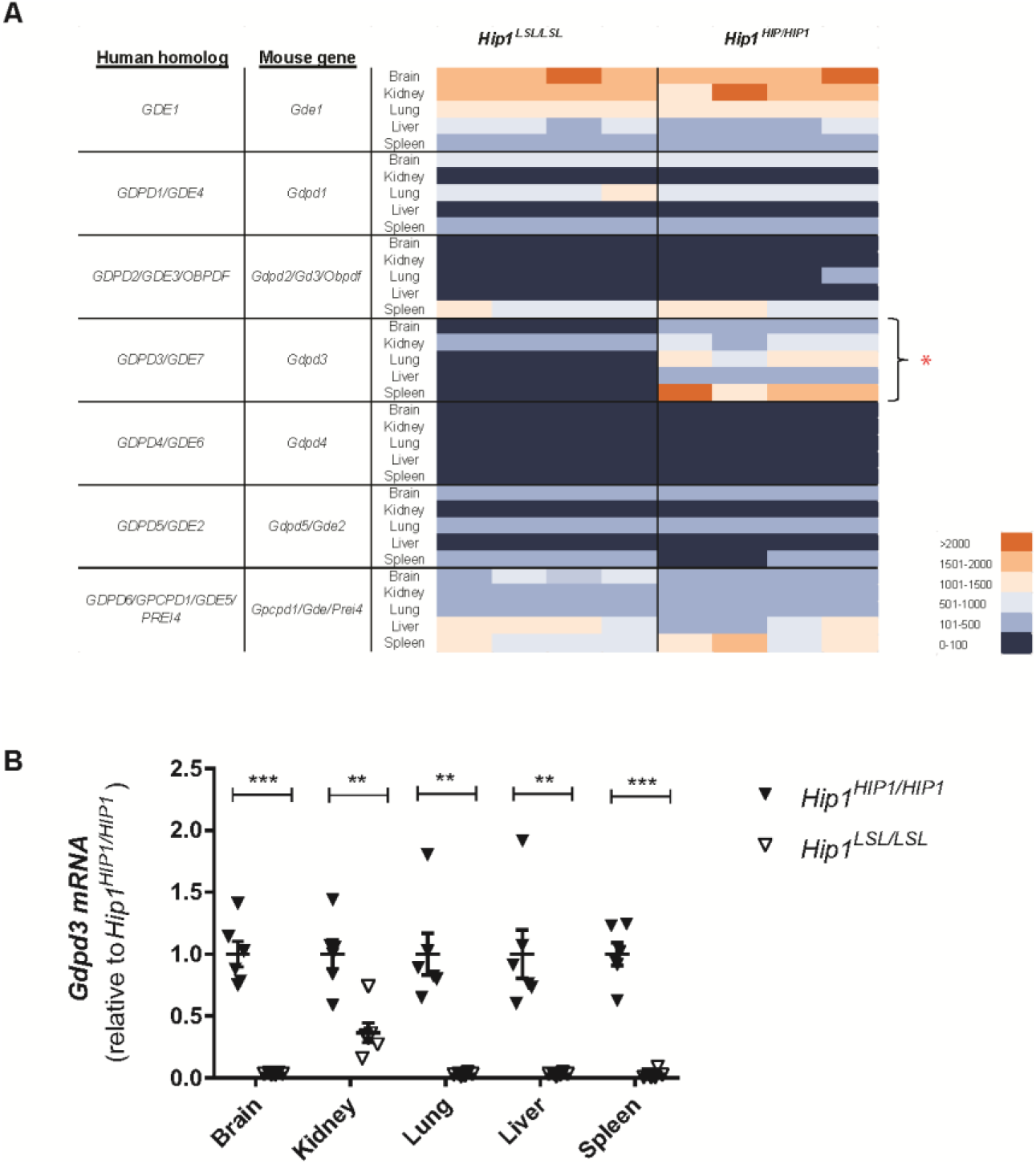
Decreased *Gdpd3* expression in *Hip1* deficient mice. **(A)** Heat map of RNA expression of the seven members of the GDPD family in *Hip1* deficient and *HIP1* rescued mice. Dark blue is the lowest and orange the highest expression. Only *Gdpd3* (*Gde7*) expression is lost in *Hip1* deficient mouse tissues (Asterisk). **(B)** Real Time PCR analysis of *Gdpd3* expression. Data were normalized to *Gapdh* and expressed as relative to the average of *Gdpd3* levels in *HIP1* rescued tissues (*Hip1^HIP1/HIP1^*). Data represent mean +/-S.E.M. n=6. p , 0.005, ***p < 0.0001 **(C)** *Gdpd3* expression in multiple tissues from *HIp1^LSL/LSL^* mice before phenotype development (1-3 months) or after phenotype development (3-7 months of age).

### Generation and analysis of *Gdpd3* knockout allele

To understand the physiological importance of Gdpd3 and to test whether Gdpd3 contributes to the Hip1 deficiency phenotypes, we generated a *Gdpd3* knockout allele by CRISPR-mediated deletion of exons 2-7 (**Figure 5A, B**). Homozygous and heterozygous knockout mice were born at predicted Mendelian frequencies and were indistinguishable at birth from wild type littermates. Levels of *Gdpd3* mRNA in lung and spleen reflected the heterozygous and homozygous state of the knockout mutation (**Figure 5C**). Heterozygous knockout tissues expressed approximately half the amount of *Gdpd3* message compared to wildtype tissue, and homozygous knockout tissue had no detectable *Gdpd3* message. In contrast, Gdpd3 deficiency did not alter *Hip1* mRNA levels indicating that *Hip1* expression is not regulated by Gdpd3.

**Figure 5.**
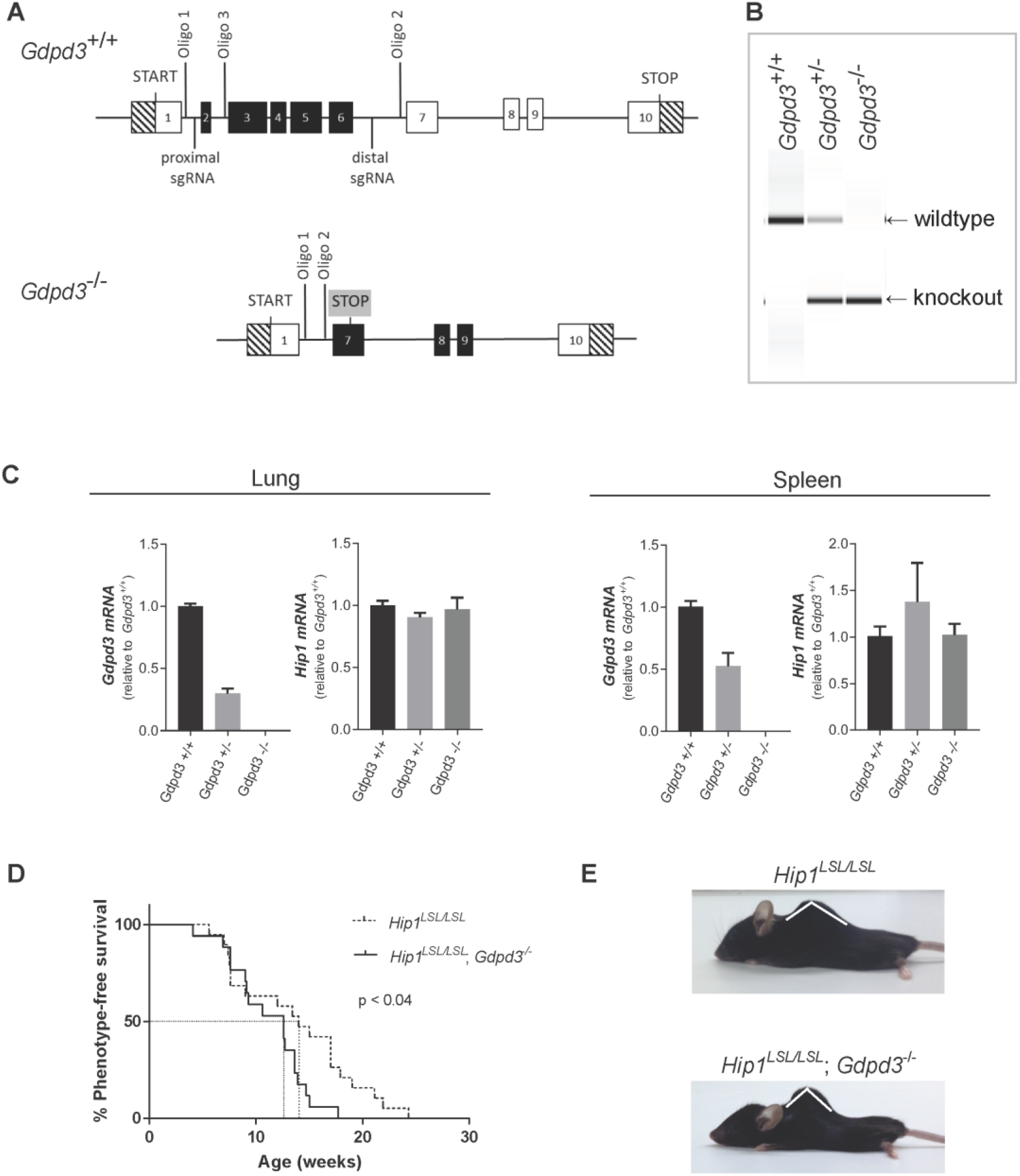
Acceleration of Kypholordosis Onset in *Hip1* and *Gdpd3* DKO mice compared to single *Hip1* knockout mice. **(A)** Schematic of the targeting strategy used to generate the *Gdpd3* knockout allele. **(B)** PCR analysis of genomic DNA isolated from tail biopsies of *Gdpd3* wild type (*Gdpd3^+/+^*), heterozygous (*Gdpd3^+/+^*), and homozygous (*Gdpd3^-/-^*) mice. Wildtype *Gdpd3* allele generates a 501bp band and the knockout allele a 290bp band with the use of oligos 1 and 3 and oligos 1 and 2 respectively. **(C)** qPCR analysis of *Gdpd3* mRNA expression levels in lung and spleen tissue of *Gdpd3^++^, Gdpd3+/* and *Gdpd3^+/^* mice. Data were normalized to *Gapdh* and expressed as relative to the average of *Gdpd3* or *Hip1* levels in *Gdpd3^+/+^* tissue. n=3 per genotype. **(D)** Kaplan-Meier curves of phenotype onset in *Hip1* knockout mice (dotted line) and *Hip1* and *Gdpd3* DKO mice (solid line). Log-rank test was used to calculate significance. Mice were phenotyped for kypholordosis as described in the methods without prior knowledge of mouse genotypes. **(E)** Representative 10 month old male *Hip1^LSL/LSL^* single knockout and *Hip1^LSL/LSL^*; *Gdpd3^-/-^* DKO mice.

Both male and female *Gdpd3* deficient mice were fertile with normal litter sizes and gender distributions. Unlike *Hip1* knockout mice who were afflicted with male infertility, as well as weight loss and a kypholordotic spinal defect by six months of age,
*Gdpd3* knockout mice displayed no consistent defects even at 2 years of age (**data not shown**). Analysis of hematoxylin and eosin stained tissues showed no abnormalities in *Gdpd3* knockout mice compared to age and gender matched controls.

### *Gdpd3* knockout mice show no changes in phosphocholine (PC)

Gdpd3 has lysophosphatidylcholine (LPC)-specific PLD (LysoPLD) activity that can generate lysophosphatidic acid (LPA) and choline (37, 38). In the first step in the biosynthesis of phosphatidylcholine, choline is phosphorylated to generate PC. We hypothesized, that the decrease in Gdpd3 in *Hip1* deficient mice could lead to the observed low PC levels (**Figure 6**). As with *Hip1* deficient tissues, we used mass spectrophotometry to profile metabolites in wild type and *Gdpd3* deficient tissues. Samples were prepared from age-and gender-matched *Gdpd3^+I+^* and *Gdpd3^+/+^* mice. To our surprise, choline and related metabolites (including PC) were at normal levels in *Gdpd3* deficient mice (**DATASET S5**). Unlike with *Hip1* deficient tissues, deletion of *Gdpd3* resulted in fewer perturbations in the metabolic profile where only a small number of metabolites showed modest changes in concentration (**DATASET S6**). It is possible that in the absence of Gdpd3, other Gdpd family members (e.g. Gdpd1 that is also thought to be a lysoPLD enzyme) compensate for its functions.

**Figure 6.**
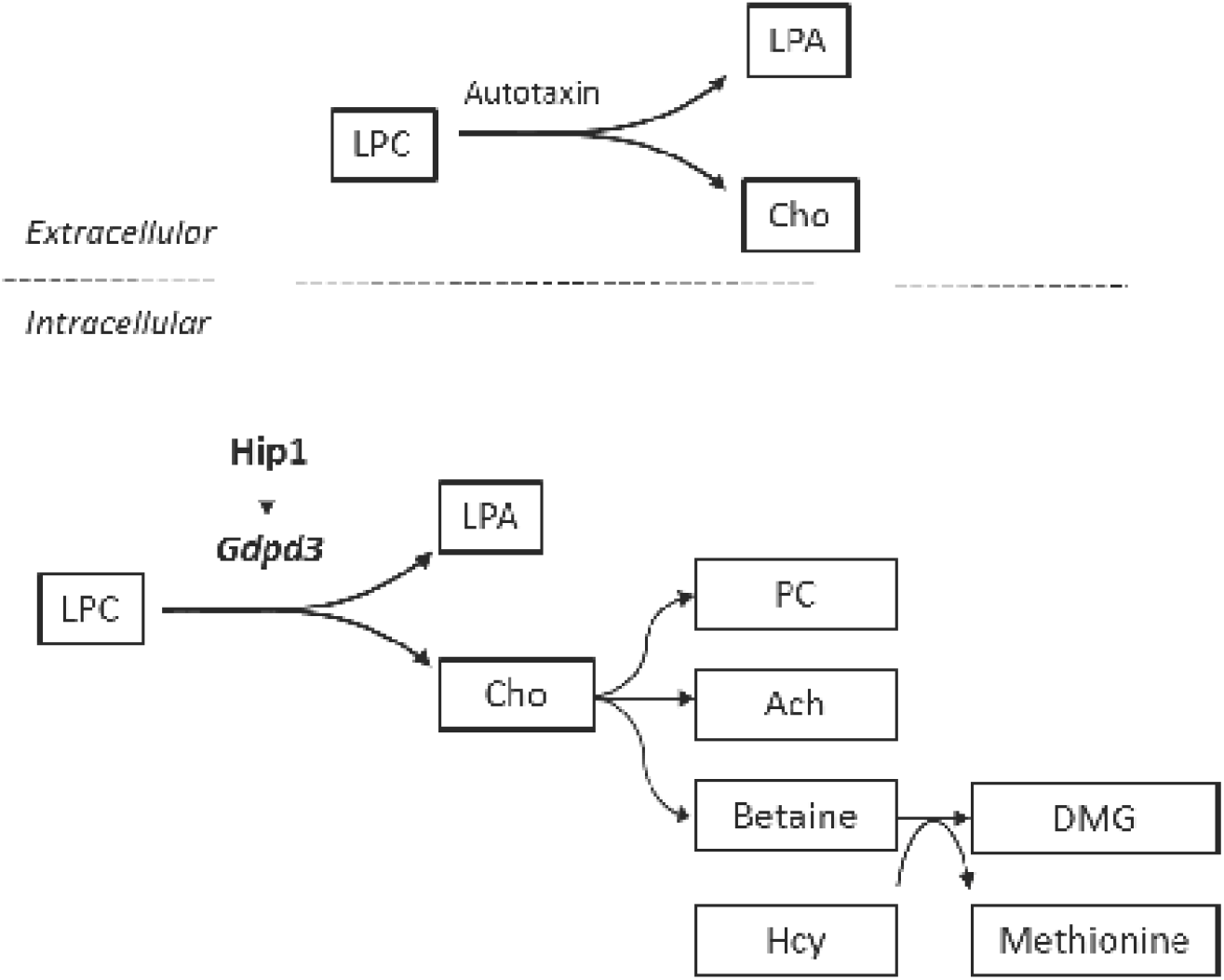
Schematic depicting a potential link between reduced Gdpd3 and alterations in choline-related metabolitesin Hip1 deficient mice. Extracellularly, LPC is metabolized to LPA and choline by the lysoPLD Autotaxin. Hip1 deficiency results in decreased Gdpd3. LysoPLD activity of Gdpd3 predicts that alterations seen in choline-related metabolites with Hip1 deficiency be due to reduced Gdpd3 levels. Changes in choline-related metabolites in liver of Hip1 deficient mice depicted. LPC=Lysophosphatidylcholione, LPA=Lysophosphatidic acid, Cho=Choline, PC=Phosphocholine, Ach=Acetylcholine, DMG=Dimethylglycine, Hcy=Homocysteine

### *Hip1/Gdpd3* double knockout (DKO) mice

*Gdpd3* deficiency hastened the onset and severity of the degenerative phenotype caused by *Hip1* deficiency. The time of onset of the kypholordosis as detected by palpation in *Hip1/Gdpd3* DKO mice was significantly earlier compared to *Hip1* single knockout mice (**Figure 5D**). All phenotyping was performed by an investigator (G.L.P.) blinded to genotype. There was no difference in the time of onset of phenotype between *Hip1^LSL/LSL^* and *Hip1*^LSLlLSL^; *Gdpd3^+I^/-* mice (**data not shown**). In a majority of cases, by the time of sacrifice due to failure to thrive, the angle of the thoracic curvature was smaller (more severe) in DKOs compared to age-and gender-matched single *Hip1* knockout mice (**Figure 5E**). These data indicate that Gdpd3 and Hip1 compensate for one another to maintain spinal homeostasis. These data also suggest that the diminished *Gdpd3* expression in the *Hip1* knockout mouse tissues (**Figure 4**) contributes to the *Hip1* knockout phenotype and the remaining small amounts of Gdpd3 provide some protection.

## DISCUSSION

The role of HIP1 in leukemia (15) and lung cancer (*20-22*) as an oncogenic fusion partner and the necessity of Hip1 for mouse homeostasis are well established (*11, 12*). Additionally, HIP1 itself is upregulated in many cancers (*24, 25, 35, 36*) and prognostic in prostate cancer (25). However, mechanisms for how HIP1 overexpression contributes to transformation (*24, 26*) and how its deficiency in mice leads to degeneration are not known. Because HIP1 binds to clathrin and AP2 (*3-6*) and alters endocytosis in cultured cells when perturbed (*40*), disrupted endocytosis that results from HIP1 abnormalities is a reasonable hypothesis for possible mechanisms to cancer, as well as mechanisms to degeneration (*41*).

*HIP1* is a large 250 kb, 32 exon gene and its second intron is a rare regulatory ATAC intron (*42*); its complex gene structure suggests that the introns and potential alternative splice products are important for maintenance of its “normal” levels. However, here we generated a *Hip1* targeted constitutive knockout with the option of conditional re-expression from a humanized allele. We showed that a single copy human *HIP1* cDNA (intron-free with the exception of intron 1), when present in all cells of the mouse (germline), was able to fully rescue *Hip1* deficiency phenotypes. Therefore, alternative splicing and most of the intervening sequences are not necessary for normal HIP1 levels and functions.

Our novel conditional *Hip1* allele allowed us to use multiple Cre alleles to express Hip1 in a tissue-specific manner to understand the role of Hip1 in spinal homeostasis. To our surprise re-expression of HIP1 specifically in the adult hematopoietic system, kidney and liver by induction of *Mx1-Cre* with pIpC was sufficient to prevent development of the *Hip1* deficiency phenotypes. Using bone marrow transplantation of Hip1 knockout bone marrow into lethally irradiated wild type mice, we found Hip1 deficiency in the bone marrow did not result in development of the Hip1 deficiency phenotype. In contrast, robust, but restricted, expression of *HIP1* in the central nervous system (both neurons and glia) using a *GFAP-Cre* transgene was insufficient to prevent the *Hip1* deficiency phenotypes. These data indicate that expression of a HIP1 cDNA in spleen, kidney and/or liver, but not in the brain, is required for mouse homeostasis.

To explore mechanisms for how Hip1 is required for homeostasis we analyzed the metabolomes and transcriptomes of *Hip1* deficient tissues. Hip1 deficiency was associated with reduced PC levels and reduced *Gdpd3* expression levels. Specifically, *Hip1* deficiency was associated with low liver, kidney and spleen PC levels. PC levels in the brain were normal. This correlates with the rescue of the *Hip1* deficiency degenerative phenotypes with HIP1 expression in adult liver, kidney and spleen, but not with HIP1 expression in the brain. The lack of PC depletion in the brain implies either that Hip1 is not involved in maintaining brain PC levels or that redundant metabolic mechanisms exist to ensure choline homeostasis in the brain. In fact, the capacity of neurons to synthesize choline *de novo* is limited and most neuronal choline comes from influx through a high-affinity choline transporter CHT1 (43). Lack of Hip1 may therefore be less likely to lead to decreased steady state PC levels in the brain compared to other tissues.

*Gdpd3* (*Gde7*) is a member of the mammalian glycerophosphodiester phosphodiesterase domain containing (Gdpd) gene family (**Figure 4A**) (*38, 39*). The common GDPD domain suggests potential functions for this family in phospholipid metabolism. However, the substrate specificity and specific enzymatic activity, as well as the physiological functions of proteins encoded by this gene family are not well understood. As recombinant proteins, both Gdpd3 (Gde7) and Gdpd1 (Gde4; Gdpd3’s closest relative) have been shown to have lysoPLD activity that catalyzes the formation of LPA and choline (*44, 45*). LysoPLD activity of Gdpd3 makes it plausible that low Gdpd3 expression in Hip1 deficient mice led to perturbations in choline-related metabolites as discussed in the results section.

The lysoPLD activity of Gdpd3 predicts that LPA may be decreased in *Hip1* deficient mice. We did, in fact, measure LPA levels in serum and tissue but found no changes with the exception of a slight elevation in *Hip1* deficient liver tissue (**data not shown**). However, serum LPA (considered tumorigenic (*46*)) is generated by Autotaxin, a PLD located on the outside surface of the cell (47). Because GDPD3 is intracellular (39), it less likely to contribute to serum LPA levels and it is not surprising that we do not see changes in serum LPA in Hip1 deficient mice (Figure 6). The high abundance of extracellularIserum LPA can complicate the measuring of tissue/intracellular LPA. Any changes in intracellular LPA levels that may occur due to loss of Gdpd3 may be confounded by serum/extracellular LPA.

Because HIP1 is expressed at high levels in many cancers (*25*) and can directly transform cells (*24*), this newly discovered alterations in choline-related metabolite levels in the *Hip1* deficient mice may not only help unravel the mechanism of degeneration associated with *Hip1* deficiency, but also help explain how HIP1 overexpression transforms cells-especially since abnormal choline phospholipid metabolism (elevated PC and diminished GPC) is an emerging hallmark of cancers (27, 48). In fact, prior published data converge on a connection between HIP1, GDPD3, cancer and choline. Specifically, *Gdpd3* has been found to be upregulated in mouse multiple myeloma (*49, 50*), HIP1 is required for growth of multiple myeloma cells (51), and transgenic overexpression of HIP1 induces myeloma like neoplasms in mice (*52*).

In addition to Gdpd3, other GDPDs have been implicated in cancers and choline pathways. For example, GDPD5 (GDE2) is elevated in breast cancer and has been correlated with elevated PC levels (53). GDPD5 has GPC-phosphodiesterase activity that generates PC. Dysregulation of choline metabolism may therefore be a shared mechanism for how altered GDPD family members could contribute to cancer development. However, each member of the GDPD family of enzymes is thought to have unique activities and unique substrate specificities; our understanding of their activities and metabolic effects will only be increased with additional, detailed *in vivo* studies.

Since HIP1 is a member of a group of endocytic oncoproteins that are hypothesized to usurp normal endocytic pathways to transform cells by increasing tumor-promoting signals (54), the decreased PC levels in the knockout mice add a new layer of complexity to the endocytosis and cancer hypothesis. By altering membrane trafficking, aberrant endocytic factors such as HIP1 could simultaneously elevate levels of several growth factor receptors by altering their endocytosis (*41, 54*), and also by influencing membrane lipid composition. It has already been shown that inhibition of endocytosis can lead to changes in membrane lipid composition (*55*), and that changes in membrane lipid composition effect endocytosis of membrane receptors (*56*). These changes in membrane lipid composition and altered receptor signaling could then influence or be influenced by choline pathway enzymes, such as Gdpd3, through feedback loops that increase PC levels in cancer and decrease PC levels in degeneration.

## Acknowledgements

We are grateful to Travis Laxson, Katherine Oravecz-Wilson, Alanna Coughran, Gunjan Singh and other members of the Ross lab for their technical assistance and intellectual contributions. We thank Xialolei Shi and Ralph Deberardinis for assistance with mass spectrometry and the Genomics Shared Resource core at the Harold C. Simmons Comprehensive Cancer Center for their expertise in RNA expression analysis. The Genomics Shared Resource is supported in part by an NCI Cancer Center Support Grant, 1P30 CA142543. This work was supported by National Cancer Institute grants to TSR (R01 CA82363-03 and R01 CA098730-01) and a Burroughs Wellcome Fund Clinical Scientist Award in Translational Research.

## Abbreviations

GDPD: Glycerophosphodiester phosphodiesterase domain
GDPD3: Glycerophosphodiester phosphodiesterase domain containing 3
HIP1: Huntingtin Interacting Protein 1
HTPIr: Huntingtin Interacting Protein 1 related
LPA: Lysophosphatidic Acid
LPC: Lysophosphatidylcholine
PC: Phosphocholine
PLD: Phospholipase D

